# A modeling-based framework to evaluate forgiveness of TB drug combinations in a BALB/c relapsing mouse model

**DOI:** 10.1101/2025.08.07.668704

**Authors:** Sylvie Sordello, Laure Brock, Alessia Tagliavini, Denise Federico, Xavier Boulenc, Marco Pergher, Emilie Huc Claustre, Darren Metcalf, Nicholas D. Walter, Gregory T. Robertson, James Clary, Alexander Berg, Khisimuzi Mdluli, David Hermann, Debra Flood, Anna M. Upton

## Abstract

Tuberculosis (TB) remains a leading cause of death due to an infectious agent. Adherence to long and complex TB treatments is supported by methods including directly observed therapy. The negative impact of missed drug doses on clinical outcomes is well-established, highlighting both the importance of adherence support and methods to quantify the ability of a regimen to continue exerting a biologic effect, during gaps in dosing known as “forgiveness” property. To explore the value of the BALB/c Relapsing Mouse Model of TB in evaluating treatment forgiveness, we assessed the impact of weekend dose holidays on the bactericidal, including RS ratio^®^, and sterilizing efficacy of RHZE/RH and BPaMZ in perspective of each drug exposure. The cure/relapse data from this study plus multiple historical studies were used to identify a nonlinear mixed-effects Emax model that was used to estimate time to cure 50% and derive time to cure 90% mice (T90). Expected time-dependent bactericidal activity and reductions in RS ratio were observed for both treatments, with more rapid decreases for the BPaMZ groups. The weekend dosing holiday significantly decreased reductions in lung CFU and RS ratio earlier in RHZE/RH treatment, but no such effect was observed for BPaMZ. Similarly, the predicted T90 was significantly greater for RHZE/RH (but not BPaMZ), with weekend doses omitted. No major drug exposure difference was observed between the 2 dosing schedules. Our results suggest BPaMZ is more forgiving of missed doses than RHZE/RH and suggests utility of this methodology to support evaluation of TB treatment forgiveness.

## INTRODUCTION

Tuberculosis (TB) was the world’s second leading cause of death from a single infectious agent in 2022, after coronavirus disease (COVID-19) and caused almost twice as many deaths as HIV/AIDS. More than 10 million people continue to fall ill with TB every year [1]. Although effective therapies are available for drug susceptible and most forms of drug-resistant tuberculosis, these are far from ideal for patients and healthcare systems. A major obstacle to the control of TB is poor adherence to lengthy, complicated and often poorly tolerated treatment regimens. Incomplete treatment can lead to increased morbidity and mortality, prolonged infectiousness and transmission, and the development of drug resistance. The impact and prevalence of missed doses of the standard of care regimen for drug susceptible TB has been clearly demonstrated through detailed meta-analyses [2]. Adherence support is therefore critical to treatment success with current TB regimens. While adherence support innovations are advancing, the current reference standard remains directly observed therapy (DOT) in which a healthcare worker observes the swallowing of the TB drugs and provides written verification. DOT is resource-intensive, presents challenges to both implementers and patients and is typically not performed during weekends. Weekend doses, therefore, cannot be confirmed and present one potential source of nonadherence. The impact of missed weekend doses – or importance of daily dosing - has been demonstrated in several analyses of clinical data [3, 4, 5]

The development of new treatment strategies could lead to shorter and better tolerated regimens. However, it is likely that new regimens will remain quite lengthy, despite the inclusion of new drugs to maximize efficacy and reduce the risk of resistance [2, 8, 9].

In 2016, the World Health Organization (WHO) developed the document Target Regimen Profiles (TRP) for TB treatments focused on shorter, less toxic and more operationally accessible regimens [10]. In 2023, an update continues to present TRP tables for the categories of RS-TB, RR-TB and Pan-TB TRP, using regimen characteristics that are largely similar with one new characteristic introduced, namely forgiveness of the regimen. Forgiveness is defined operationally as “the degree to which regimen efficacy is unaffected by suboptimal adherence”, highlighting the importance of the relationship of treatment adherence to its efficacy. Two factors are thought to influence forgiveness: (1) the pharmacokinetic (PK) profile of drugs in the regimen (*i*.*e*., how long drugs remain at or above inhibitory concentrations in tissue) and (2) post-antibiotic effect (PAE) which is the lag after drug exposure before bacteria recover and resume growth (*i*.*e*., after drugs are entirely cleared) [11, 12].

Forgiveness of TB drug regimens is difficult to explore in humans for ethical, technical, and clinical reasons. This creates several concerns. First, the drug composition of regimens, due to differences in mechanisms of action, as well as in dosing, can alter forgiveness for regimens of different lengths [10]. Further it is plausible that the consequence of missed doses will be amplified in shorter treatments versus longer treatment regimens, as there are fewer doses given during the treatment interval. The WHO TRP statement recognized that forgiveness is difficult to measure and encouraged assessment of forgiveness in preclinical models.

To respond to this need, we assessed two clinically-relevant reference regimens, the existing standard drug susceptible TB regimen (RHZE/RH) and the shorter and more potent SimpliciTB regimen (BPaMZ), in the conventional BALB/c subacute TB infection mouse model [13].

Murine models of TB have been used for more than 50 years for evaluation of new TB drugs and regimens [14] and are well established as preclinical models of disease. The immunocompetent BALB/c mouse model of TB has been used extensively to evaluate the relationship between bactericidal and sterilizing efficacy and treatment duration for TB drug regimens. Endpoints typically include the bacterial burden in mouse lung at the end of treatment (end of treatment colony forming unit [CFU]), compared to pre-treatment lung CFU to assess bactericidal activity over time, and, to assess sterilizing activity as the proportion of mice “relapsing” after each treatment duration, defined as the presence of any lung CFU following a period (usually 12 weeks) off treatment [15, 16, 17, 18, 19 and 20]. A mouse model of TB when used with a “relapse” endpoint is referred to as the Relapsing Mouse Model (RMM) of TB.

Evaluation of sterilizing efficacy of a drug regimen in this model of TB historically utilized conventional group-to-group statistical comparisons within a given study, designed to evaluate the proportion of mice relapsing at a relatively small number of treatment durations (e.g., three to five). Lenaerts et al. [19] demonstrated that this approach requires at least 15 animals per regimen to detect a 50% reduction in relapse probability at a given treatment duration with at least 80% power. This analytical strategy based upon a point-by-point comparison of relapsing proportions limits the utility of the model as the differences between candidate regimens are often much smaller. Further, the focus on comparative efficacy at prespecified treatment durations requires careful selection of multiple treatment durations (and larger numbers of mice) to be tested to ensure identification of regimens that achieve significant rates of cure (e.g., 10% or lower probability of relapse) with a shorter treatment duration than the current standard of care regimen. Further, results from RMM studies have typically been reported with limited raw data to support statistical analyses, therefore limiting comparisons, interpretation, and decision-making across RMM studies and regimens.

In recent years, innovations in both experimental design and data analysis have enabled improved evaluations of the relationship between cure and treatment time for TB drug regimens both within and across studies, as well as reduction in animal use incurred by these studies. First, Mourik and colleagues introduced experimental designs that used smaller groups of mice coupled with more frequent and numerous timepoints, with mathematical modeling of the data to assess the relationship between relapse/cure and treatment duration [20]. This group also advocated “bundling” of RMM data across studies to better compare tested TB regimens to support translation [21]. More recently, Berg et al [22] showed through the application of model based meta-analysis approaches to a large data set of 28 studies, that the understanding and interpretability of regimen performance in the RMM was improved. An existing published model was developed by Berg et al. [22] using a logistic regression model to characterize the treatment duration-dependent probability of relapse for each regimen and identify relevant covariates contributing to inter-study variability.

Given the advantages of model-based meta-analysis in enabling cross-study comparisons, we have adopted this approach, coupled with experimental designs inspired by Mourik et al, for evaluation of the treatment duration-cure relationship for TB drug regimens.

An alternative measure of treatment effect is a novel pharmacodynamic (PD) marker called the RS ratio. Unlike culture, which estimates bacterial burden, the RS ratio quantifies *Mtb* rRNA synthesis, thereby providing a measure of bacterial activity, that indicates the degree of ongoing *Mtb* rRNA synthesis in the lungs of *Mtb*-infected mice [23, 24, 25].

Since a large bacterial population may have an inactive phenotype or a small bacterial population may have an active phenotype, the RS ratio provides information distinct from culture. The speed and degree to which regimens suppress the RS ratio in mice appears to correlate with their treatment-shortening activity in mice [23].

This work aims to explore the value of the BALB/c RMM of TB, used with culture-based and RS ratio endpoints, within a modeling-based framework, for evaluation of forgiveness of TB drug regimens. Weekend dosing holidays in mice are used to simulate missed weekly doses in patients. We also introduce a modeling and simulations approach to design of RMM studies with optimal animal use, and a nonlinear mixed-effects Emax model based on historical data and data generated via this study to estimate time to 50% cure and derive time to 90% cure (T90). Steady-state plasma exposures as well as lung (supplementary data) exposures were also assessed to evaluate the potential of drug exposure-related causes of differences in efficacy between the same regimen dosed 5 of 7 (5/7) and 7 of 7 (7/7) days per week.

The two clinically-relevant reference regimens selected, the existing standard drug susceptible TB The development of new treatment strategies could lead to shorter and The development of new treatment strategies could lead to shorter and The development of new treatment strategies could lead to shorter and The development of new treatment strategies could lead to shorter and The development of new treatment strategies could lead to shorter and The development of new treatment strategies could lead to shorter and The development of new treatment strategies could lead to shorter and The development of new treatment strategies could lead to shorter and The development of new treatment strategies could lead to shorter and The development of new treatment strategies could lead to shorter and The development of new treatment strategies could lead to shorter and The development of new treatment strategies could lead to shorter and The development of new treatment strategies could lead to shorter and The development of new treatment strategies could lead to shorter and The development of new treatment strategies could lead to shorter and The development of new treatment strategies could lead to shorter and regimen (RHZE/RH) and the shorter and more potent SimpliciTB regimen (BPaMZ), are composed of diverse drugs with respect to both modes of action and pharmacokinetics (PK). In an optimal study design, we compared conventional and novel pharmacodynamic (PD) markers and time to cure (relapse data modelling) among mice allowed a 2-day treatment interruption each week (5/7) versus those with uninterrupted daily treatment (7/7).

## RESULTS

### Study design optimization using a modelling and simulations approach

Following the principles described in Mourik et al [19] and utilizing the curated data set and model established in Berg et al [22], simulations were performed to generate study design options for the evaluation of RHZE/RH and BPaMZ, dosed either 5/7 or 7/7 in the BALB/c RMM, that minimize animal use while seeking to maximizing the utility of relapse data in predicting time to cure in 90% of mice (T90) for each drug combination and treatment regimen. A simulation study was performed to investigate the relative performance of proposed RMM study designs based on key performance parameters (i.e. bias and precision of model-based estimates obtained from various study designs). Simulations were performed in accordance with the methodologies described in Clary et al (to be submitted to AAC) [26]. The simulated data was combined with relapse historical data and re-estimated to determine the ability of the design to estimate T50 and derive T90 with minimal bias compared to simulated value and minimal number of mice needed per timepoint. Based on these criteria, the design for the relapse portion of the study was optimized and selected (represented in Figure 1, Table 1). In addition, to enable assessment of bactericidal activity over time for each combination and treatment regimen, groups were added to allow for lung bacterial load enumeration directly after completion of each treatment duration. In addition, changes in RS ratio biomarker with each combination/regimen were assessed. Following 8 weeks treatment a pharmacokinetics sampling schedule was integrated in order to assess related drug exposures at steady-state, as well.

**Table 1.** Treatment groups: number of mice per treatment group at each endpoint for each readout during the different phase of the study: Each phase, infection (day post infection: dpi) treatment (Tx) and Relapse (R) duration are indicated in week (W). the number of mice used per readout is indicated before the parameter measured in bracket (CFU, RS for RS ratio or PK). For the relapse only the number of mice used per time point is indicated for measuring the bacterial load at the end of 12 weeks off treatment (+12WR)

|  | Infection phase |  | Treatment phase (Tx) |  |  |  |  |  |  |  |  |  |  |  |  |  |  |  |
| --- | --- | --- | --- | --- | --- | --- | --- | --- | --- | --- | --- | --- | --- | --- | --- | --- | --- | --- |
| Treatment | Day -13<br>(1 dpi) | Day 0<br>(14 dpi) | end of<br>2W Tx | end of<br>4W Tx | end of<br>6W Tx | end of<br>8W Tx | end of<br>10W<br>Tx | end of<br>12W<br>Tx | end of<br>14W<br>Tx | end of<br>16W<br>Tx | 2W<br>Tx +<br>12W<br>R | 4W<br>Tx +<br>12W<br>R | 6W<br>Tx +<br>12W<br>R | 8W<br>Tx +<br>12W<br>R | 10W<br>Tx +<br>12W<br>R | 12W<br>Tx +<br>12W<br>R | 14W<br>Tx +<br>12W<br>R | 16W<br>Tx +<br>12W<br>R |
| Untreated | 5(CFU) | 5(CFU)<br>+ 3(RS) | 5 (RS) | - | - | - | - | - | - | - | - | - | - | - | - | - | - | - |
| BPamZ 7/7 | - | - | 5<br>(CFU)<br>+ 5(RS) | 5(CFU)<br>+ 5(RS) | 5(CFU)<br>+ 5(RS) | 5(CFU) +<br>3(RS) +<br>6(PK) | - | - | - | - | 6 | 6 | 6 | 6 | - | - | - | - |
| BPamZ 5/7 | - | - | 5(CFU)<br>+ 5(RS) | 5(CFU)<br>+ 5(RS) | 5(CFU)<br>+ 5(RS) | 5(CFU) +<br>3(RS) +<br>6(PK) | 5(CFU) | 5(CFU) | - | - | 6 | 6 | 6 | 6 | 6 | - | - | - |
| 2RHZE/<br>2RH 7/7 | - | - | 5(CFU)<br>+ 5(RS) | 5(CFU)<br>+ 5(RS) | 5(CFU)<br>+ 5(RS) | 5(CFU) +<br>3(RS) +<br>6(PK) | 5(CFU) | 5(CFU) | 5(CFU) | 5(CFU) | - | - | - | 6 | 6 | 6 | 6 | 6 |
| 2RHZE/<br>2RH 5/7 | - | - | 5(CFU)<br>+ 5(RS) | 5(CFU)<br>+ 5(RS) | 5(CFU)<br>+ 5(RS) | 5(CFU) +<br>3(RS) +<br>6(PK) | 5(CFU) | 5(CFU) | 5(CFU) | 5(CFU) | - | - | - | 6 | 6 | 6 | 6 | 6 |

**Figure 1.**
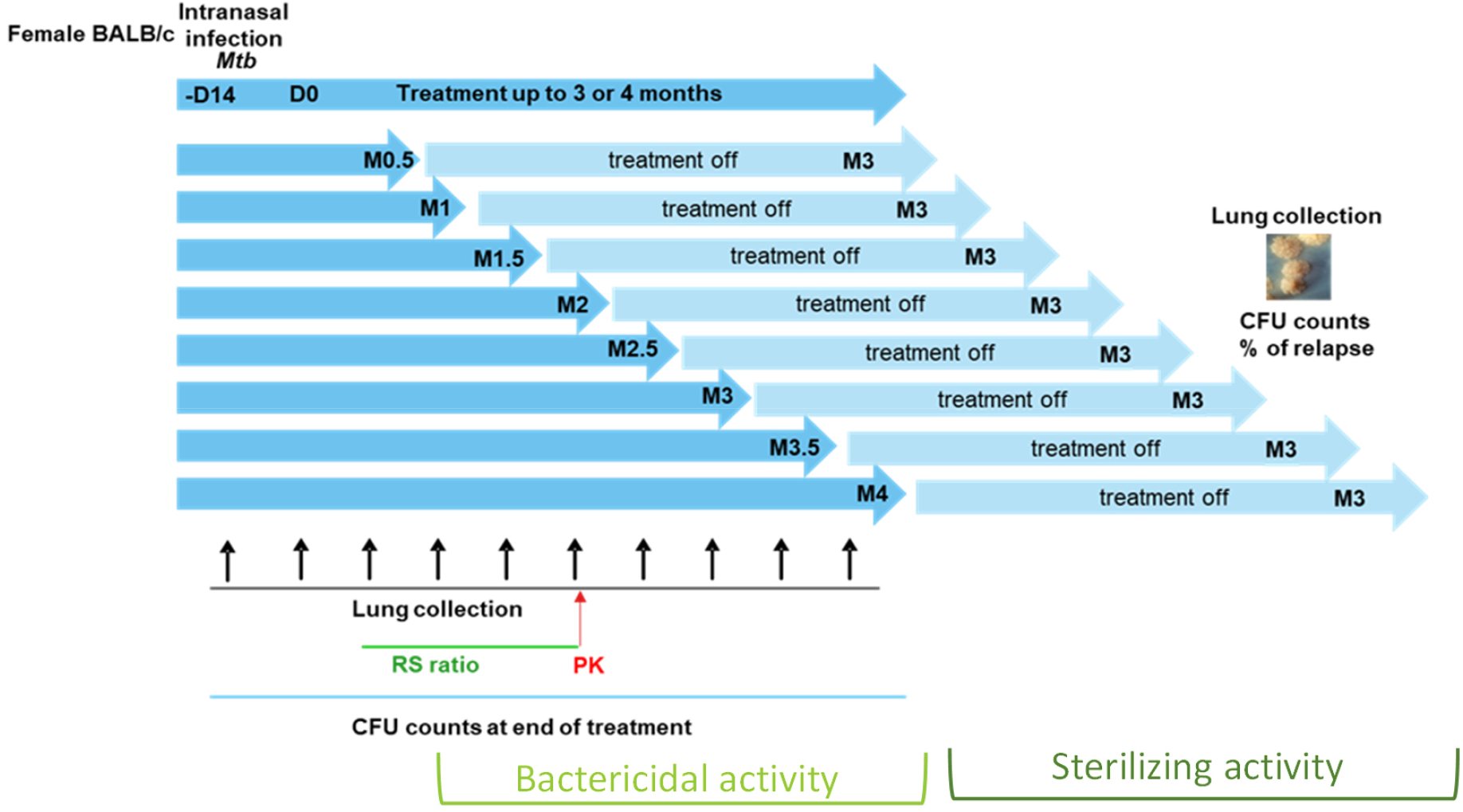
RMM Study Design

### Impact of 5/7 versus 7/7 dosing on bactericidal and on RS ratio activity

Following intranasal infection of BALB/c mice with *M. tuberculosis* H37Rv, the bacterial burdens in the lung at 1 day (Day-13) and 14 days (Day 0) post infection were 4.5 +/-0.09 and 7.28 +/-0.26 Log10 CFU/lung respectively (mean +/-sd). The drug combinations, BPaMZ and RHZE/RH were dosed daily per oral gavage either 5/7 or 7/7 days per week. The 4-drug combination RHZE was dosed for 2 months, followed by dosing with only RH for an additional 2 months (2RHZE/2RH). Mice were dosed for 2, 4, 6, 8, 10, 12, 14 or 16 weeks. As anticipated based on prior work [13, 14, 15, 16, 17 and 18] both combinations displayed significant bactericidal activity over time, reducing CFU in lung at the end of treatment, compared to the level at Day 0 (start of treatment).

Bacterial burdens in lungs declined significantly for all treatment groups starting from the 2-week treatment timepoint. From the 4-week treatment time points onward, some mice displayed culture negativity in the BPaMZ 5/7 and 7/7 dosing groups (Figure 2). BPaMZ had greater bactericidal than 2RHZE/2RH activity, achieving culture negativity in all mice at week 8 (Figure 2). The RS ratio also declined significantly faster and a more profound reduction was seen for BPaMZ than for 2RHZE/2RH (Figure 3), as reported previously [23].

**Figure 2.**
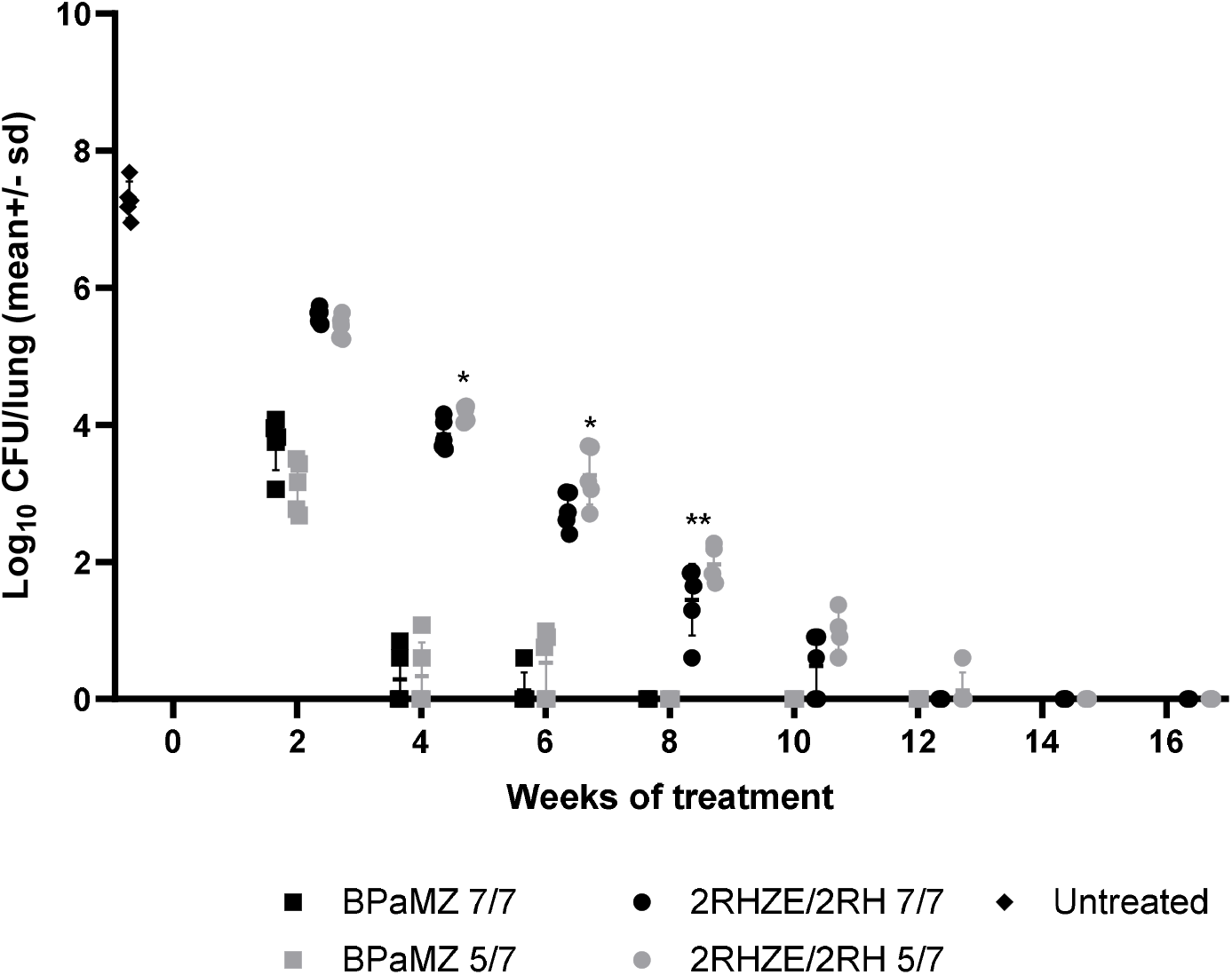
Mouse lung CFU following treatment with RHZE/RH and BPaMZ dosed 5/7 and 7/7 days per week. Whole lung CFU of BALB/c mice, intranasally infected with *M*.*tb* H37Rv, after different durations of oral treatment with RHZE/RH or BPaMZ, dosed 5/7 or 7/7. Treatments were initiated 2-weeks post infection. Statistical test results for dosing days comparison on Log10 CFU/lung. *: p<0.05; **: p<0.01.

**Figure 3.**
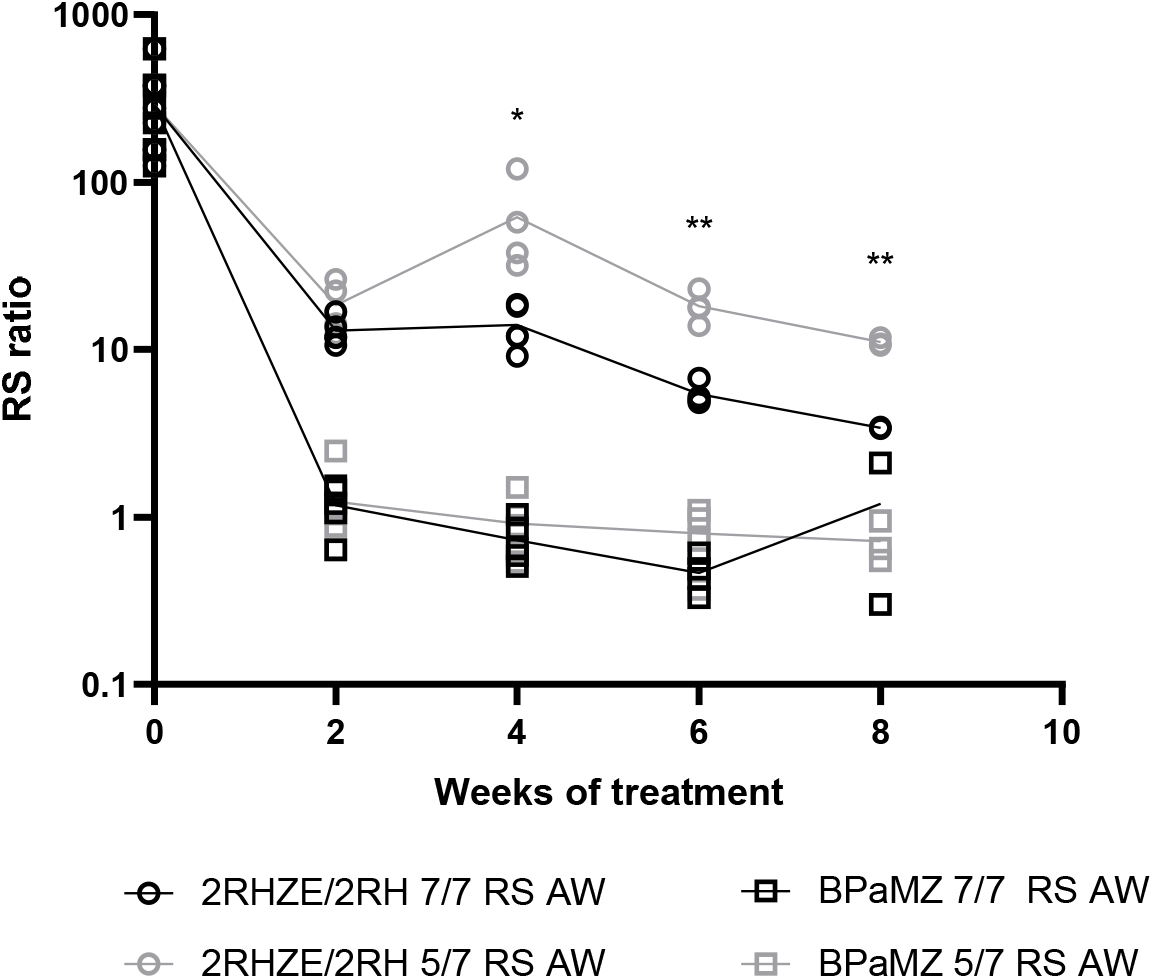
RS ratio in Mouse Lungs, following treatment with RHZE/RH and BPaMZ dosed 5/7 and 7/7 days per week. RS ratio® in lungs of BALB/c mice, intranasally infected with *M*.*tb* H37Rv, after different durations of oral treatment with RHZE/RH or BPaMZ, dosed 5/7 or 7/7. Treatments were initiated 2-weeks post infection. At different time points, lungs were collected for RS ratio® quantification. Analysis was performed on 1/3 lung. Statistical test results for dosing days comparison on RS ratio. *: p<0.05; **: p<0.01.

No significant differences were observed in the overall time to culture negativity in mice given BPaMZ 5/7 or 7/7 days per week. In contrast, 12 or 14 weeks of 2RHZE/2RH treatment were required to achieve negative lung cultures following the daily 7/7 or 5/7 dosing, respectively. In addition, for 2RHZE/2RH dosed daily (7/7), mice had a significantly lower lung CFU burden at 4-, 6- and 8-weeks of treatment than mice allowed weekend drug holidays (5/7) (Figure 2), as reported previously [18]. There was no significant CFU difference between 7/7 and 5/7 dosing at weeks 2 and 10. Similarly, mice dosed with 2RHZE/2RH daily had significantly greater suppression of the RS ratio at weeks 4, 6 and 8 relative to mice allowed weekend holidays (Figure 3), while no such difference was observed in the BPaMZ-treated mice either for CFU (Figure 2) or RS ratio (Figure 3).

### Change in CFU and RS ratio during a short treatment interruption

In order to explore the impact of a short interruption of treatment on the PD parameters evaluated in this study, we compared both lung CFU and RS ratio in lungs collected either 24h or 96h post last dosing following 2 weeks of treatment. For both RHZE and BPaMZ, there was no significant difference between CFU burdens with a 24h (1-day) or 96h (4-day) recovery period after treatment cessation (Figure 4A). For RHZE, the RS ratio increased significantly during this short period of time following end of treatment consistent with physiologic recovery following treatment. Specifically, between post-treatment days one and four, the median RS ratio rose from about 17 to 75 with 5/7 dosing and rose from 12 to 80 with 7/7 dosing (Figure 4B). By contrast, for BPaMZ, there was no significant change in the RS ratio between post-treatment days one and four.

**Figure 4.**
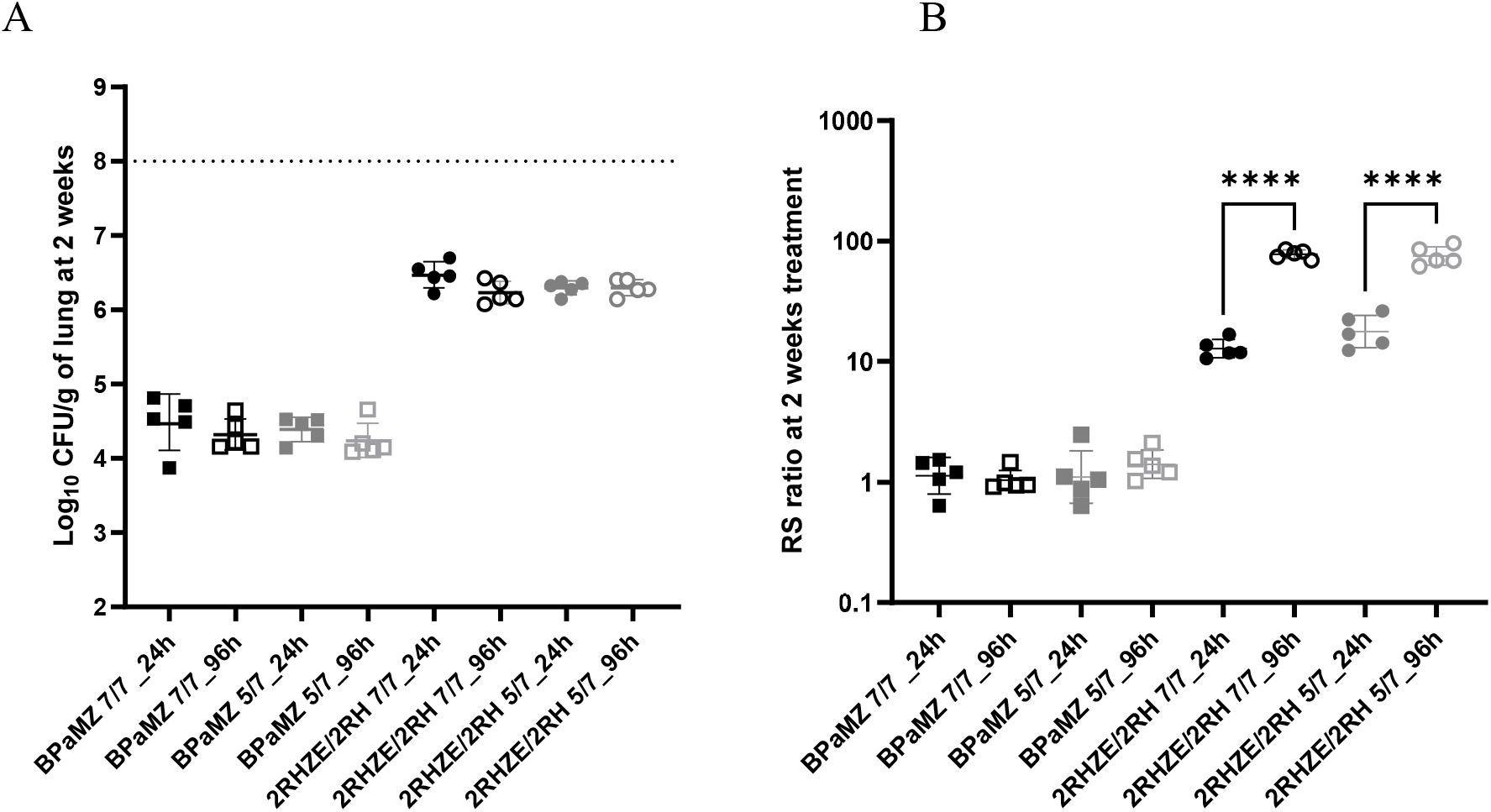
Mouse lung CFU (A) and RS ratio (B) measured 24h or 96h post last dosing at the end of 2 weeks of treatment. Lungs of BALB/c mice, intranasally infected with *M*.*tb* H37Rv, after 2 weeks treatment durations of oral treatment with RHZE/RH or BPaMZ, dosed 5/7 or 7/7 initiated 2-weeks post infection, were split in 2/3 for CFU analysis and 1/3 for RS ratio analysis. Lungs were collected 24h or 96h post last dosing. Statistical test results for dosing days comparison on Log10 CFU/lung. ****: p<0.0001.

### Significant increase in time to 90% cure (T90), derived using an Emax model, for 2RHZE/2RH but not BPaMZ, with a weekend dosing holiday

To establish the relationship between the tested treatment regimens and time for treatment required to achieve durable cure in mice (T90), we quantified the proportion of mice that were culture positive (i.e. relapse) twelve weeks after the end of treatment (i.e., 2, 4, 6, 8, 10, 12, 14, or 16 weeks treatment durations respectively). For BPaMZ-treated groups, no or a very low number of mice relapsed following 4 weeks of treatment, whereas 100% of mice relapsed following 2 weeks of treatment (Table 2). No relapse events were recorded for any BPaMZ-treated mice after 6-weeks or more of treatment (Table 2). In contrast, it took at least 16 weeks of treatment with 2RHZE/2RH to achieve 100% cure (no relapse) or appreciably low proportions of relapse events in mice (Table 2).

**Table 2.** Percentage of mice with evidence of relapse (absence of cure) for each combination (Positive culture mice*/total number of mice)

| Drug Regimen | Weeks of Treatment Prior to the 12-week Post-Treatment Relapse Phase |  |  |  |  |  |  |  |
| --- | --- | --- | --- | --- | --- | --- | --- | --- |
|  | 2W | 4W | 6W | 8W | 10W | 12W | 14W | 16W |
| BPamZ 7/7 | 100%<br>(6/6) | 0%<br>(0/6) | 0%<br>(0/6) | 0%<br>(0/5) | - | - | - | - |
| BPamZ 5/7 | 100%<br>(6/6) | 16.6%<br>(1/6) | 0%<br>(0/6) | 0%<br>(0/6) | 0%<br>(0/6) | - | - | - |
| 2RHZE/2RH 7/7 | - | - | - | 100%<br>(6/6) | 66.6%<br>(4/6) | 0%<br>(0/6) | 16.6%<br>(1/6) | 0%<br>(0/6) |
| 2RHZE/2RH 5/7 | - | - | - | 100%<br>(6/6) | 100%<br>(6/6) | 50%<br>(3/6) | 33.3%<br>(2/6) | 16.6%<br>(1/6) |

Modeling of these relapse data was conducted to estimate time to 50% cure (T50) and derive time to 90% cure (T90) per combination and treatment regimen. Model-based analysis was performed using a nonlinear mixed-effects modelling approach, wherein a logistic Emax model was developed based on observed percentage of cure/relapse data obtained pooling together historical data with data of this study. Estimations of T50 parameters were made, as well as derivation of T90 values for each treatment and dosing schedule (Figure 5). For BPaMZ and 2RHZE/2RH dosed 5/7, derived T90s were 1.29 and 4.03 months respectively. For 7/7 dosing regimens, BPaMZ and 2RHZE/2RH had derived T90s of 1.00 and 3.30 months, respectively (Table 3). Taken together, this indicates that the effect of the weekend dosing holiday significantly extended T90 by 0.7 months, compared to the daily-dosed regimens, for 2RHZE/2RH (p= 0.025), but had no significant impact on BPaMZ (p=0.110), where the shift of T90 was about 0.3 months only (Table 3).

**Table 3.**
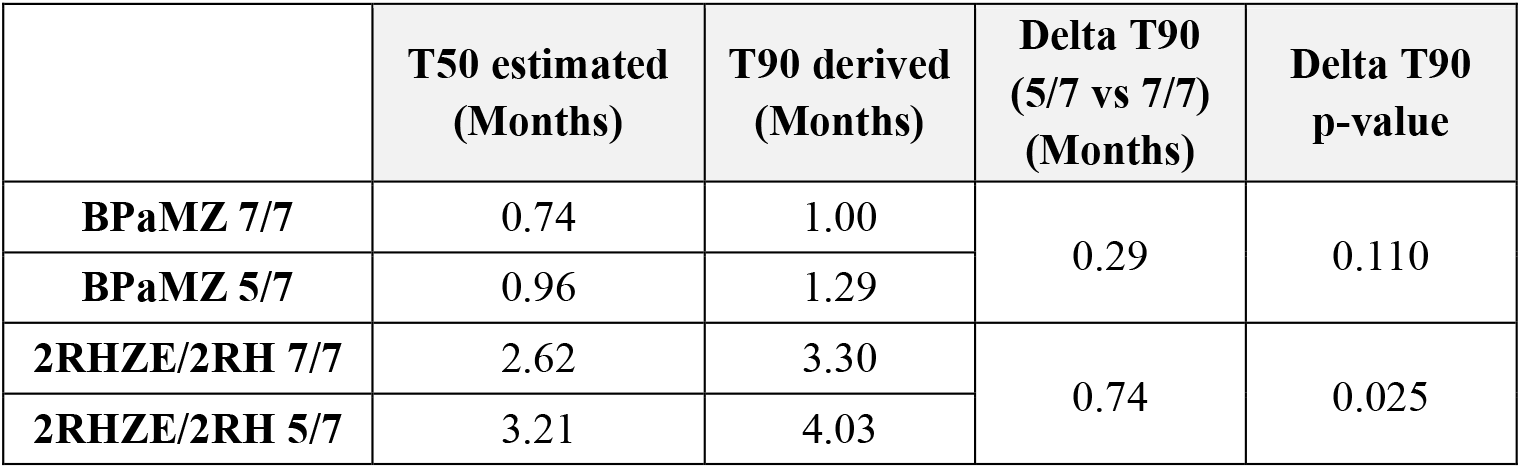
Estimated T50 and Derived T90 Values for Test Combinations and Regimens.

**Figure 5.**
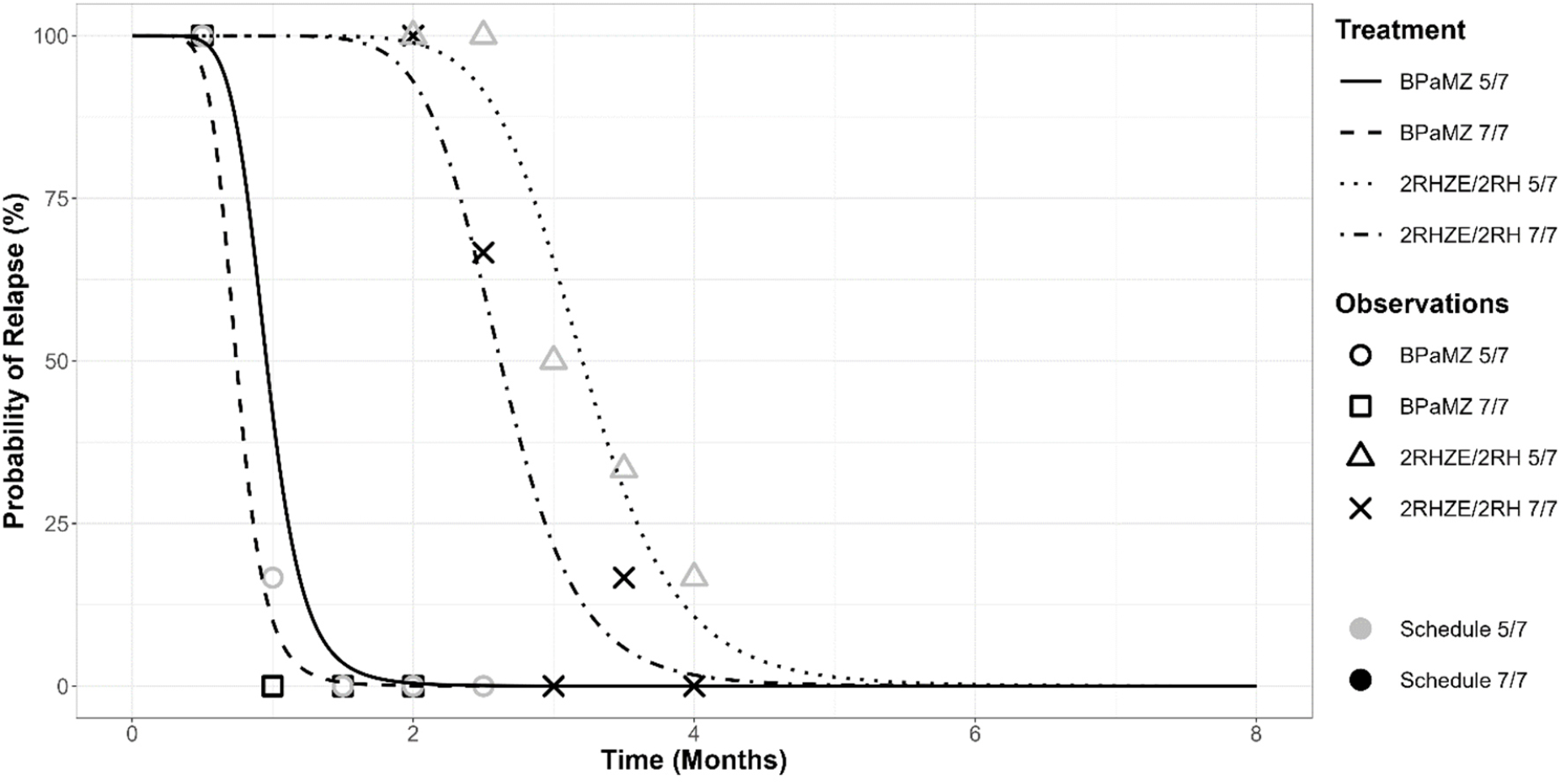
Probability of Relapse with Treatment Duration, for BALB/c Mice treated with RHZE/RH and BPaMZ. Sterilization curves indicating the probability of relapse over treatment time, constructed by fitting observed relapse data (Table 2) to an Emax model developed using a large historical RMM dataset. Observed relapse data are indicated for each test regimen using open symbols and crosses. The time to 50% cure/relapse is estimated from these curves, and the time to 90% cure is derived utilizing time to 50% cure estimates together with steepness of the curve (gamma) as explained in Methods.

### Limited impact of dosing 5/7 versus 7/7 days per week on drug exposures is only evident for pretomanid and moxifloxacin

We assessed drug exposures achieved during the RMM study, to evaluate the impact of the altered dosing schedules on overall drug levels at steady state. Following 8 weeks of treatment, evaluation of blood PK was performed, following the 5^th^ day of dosing. Found drug levels in blood were generally consistent with literature values [26, 27, 28, 29, 30]. Similar exposures were observed for drugs given 5/7 and 7/7 days per week, with the exception of pretomanid and moxifloxacin. Although Pa and M were still detectable 24h after the previous administration (i.e. Ctrough > LOQ supplementary data) when administered 7/7 (Figure 6) both drugs were below the LOQ when administered 5/7. However, even for these two drugs, the post administration concentrations in blood were similar between the two dosing schedules, with a minor impact of the low Ctrough concentrations detected after the 7/7 regimen.

**Figure 6.**
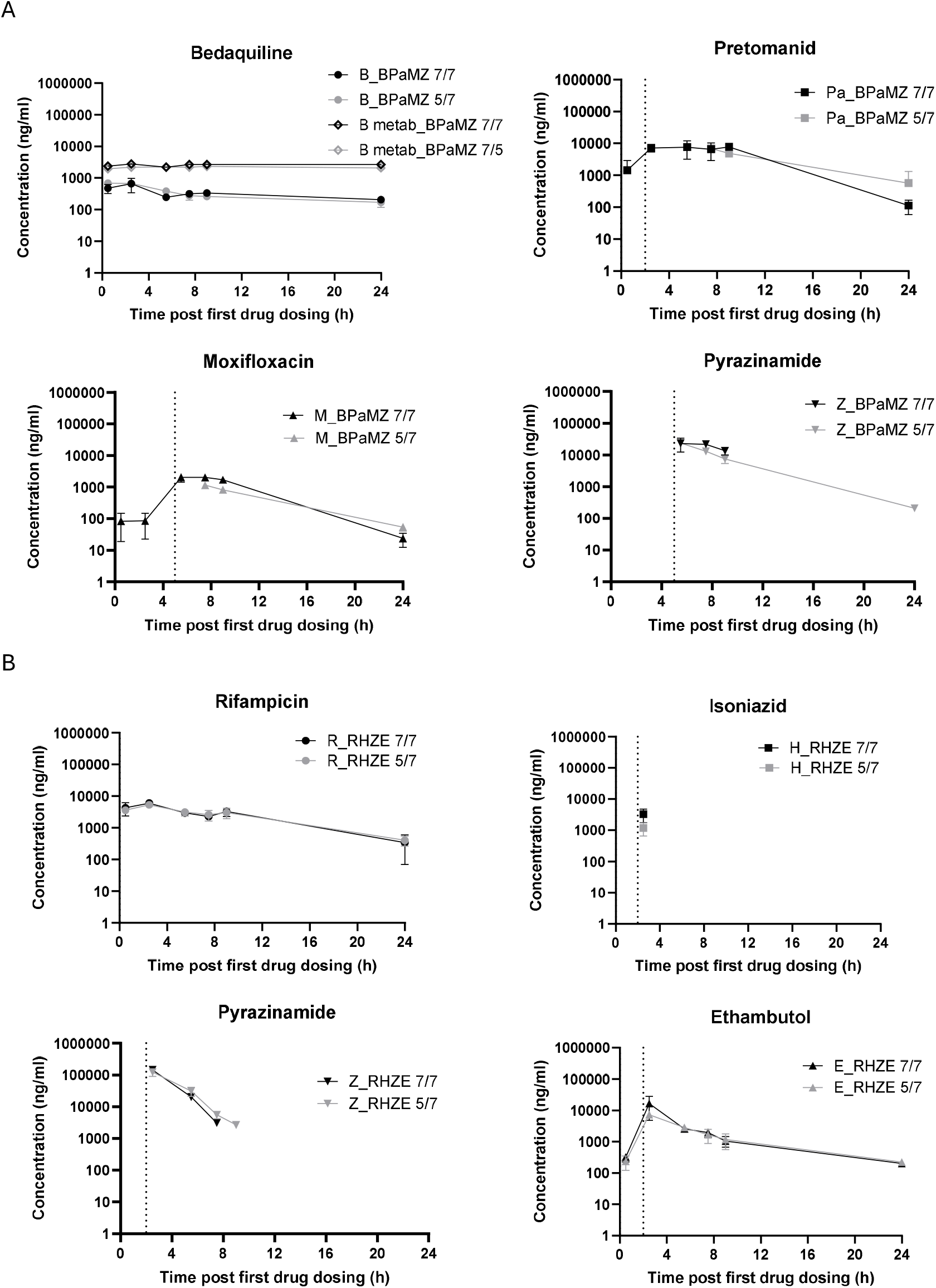
Individual blood PK after 8 weeks of treatment on the fifth day of the last week of treatment. After 8 weeks of treatment on the 5^th^ day of the last week, for each combination (6 mice per group) blood was collected at 0.5, 2.5, 5.5, 7.5, 9 and 24 hours post-first compound dosing. Individual drug exposure was determined in BPaMZ (A) and RHZE (B) after 5/7 or 7/7 dosing. Of note: Dotted Line in graph corresponds to time of dosing of corresponding drug.

## DISCUSSION

The optimized study design described herein, supported testing the impact of missed TB drug doses – simulated using a weekend dosing holiday in mice-on standard and novel pharmacodynamic parameters and overall steady-state drug exposures. Key parameters included, change in CFU lung burdens and RS ratio at the end of treatment, as well as proportion of mice undergoing relapse 12 weeks following different time of treatment completion, which were used to derive time to achieve 90% of cure (T90) for two clinically relevant benchmark drug regimens, 2RHZE/2RH and BPaMZ, given 5/7 (weekend holiday) and 7/7 (daily dosing).

Building on the prior work of Mourik et al, and Berg et al, we made use of modeling approaches both in terms of both study design and data analysis to gain a greater understanding of the relationship between treatment duration and relapse probability, while minimizing animal numbers and enabling future cross-study comparisons. Data from this study will be submitted to the C-PATH APEX platform (https://c-path.org/tools-platforms/tb-apex) to facilitate similar approaches by other research groups with an overarching aim to share data towards minimizing animal use while improving understanding of the performance of TB drug treatments in the RMM.

The absolute and relative time-dependent bactericidal activity of the 2RHZE/2RH and BPaMZ regimens given 5/7 days per week was consistent with previous published data [15, 16, 17, 18 and 19]. Similarly, the impact of 2RHZE/2RH and BPaMZ treatment on RS ratio was consistent with prior reports [23 and 24] with a larger decrease in RS ratio observed from 2 weeks onwards, for the BPaMZ groups compared to the 2RHZE/2RH groups.

Although derived via differing methodology, the derived T90s of 1.29 and 4.03 months for the 2RHZE/2RH and BPaMZ given 5/7 days per weeks are similar to published values [20, 21].

When the two drug combinations were dosed 7 days per week, the derived T90 values were reduced by around 20% for each combination. The difference observed between T90s achieved following daily dosing (7/7) and when weekend dosing holidays were observed (5/7, simulating missed doses), represents a delay to achieve 90% cure by about 1.3 weeks for BPaMZ and 3.2 weeks for 2RHZE/2RH. This difference in T90 for 5/7 versus 7/7 dosing is statistically significant for 2RHZE/2RH only. To our knowledge, this is the first study reporting sterilizing treatment outcomes when TB drug regimens have been evaluated following daily dosing 7 days per week in a relapsing mouse model of TB.

The significant differences in both kinetics of bactericidal responses (measured by lung CFU burdens reduction) and reduction in RS ratio during the first 8 weeks of treatment when 2RHZE/RH was dosed 5/7 days per week versus 7/7 daily dosing, were associated with a significant increase in the T90 in mice. No such differences were observed for BPaMZ, regardless of the dosing schedule.

Our evaluation of a brief treatment interruption after 2 weeks of treatment confirmed previous observations that the RS ratio “rebounds” when RHZE is stopped suggesting the *Mtb* begin “physiologic recovery” that is not observable via assessment of CFU [32]. Importantly, expanding on these previous findings, we demonstrate here for the first time that the RS ratio did not rebound after treatment with BPaMZ. This is consistent with the apparent forgiveness of BPaMZ and might be attributable to either the long half-life of bedaquiline (*i*.*e*., PK) [28, 29, 34] or to the distinct pattern of physiologic injury and adaptation caused by bedaquline-based regimens (*i*.*e*., PAE) [33].

Based on the data reported herein, differences in drug exposures between 2RHZE/2RH given 5/7 versus 7/7 days per week, do not readily explain the impact of the weekend holiday on treatment outcomes. The exposures of all drugs in this combination were similar at the time points evaluated (after the 5^th^ dose given in the 8^th^ week of dosing) which is assumed to represent steady state. However, we cannot rule out differences in exposure dynamics that may have occurred during weekend breaks, or earlier in the course treatment. On the other hand, modest differences were observed for exposures of pretomanid and moxifloxacin between 5/7 and 7/7 dosing schedules, but this did not have a significant impact on the bactericidal activity, RS ratio reduction over time, or the T90 for BPaMZ. As for 2RHZE/2RH, exposure dynamics over treatment time are not known nor PK during weekends and it is possible that drugs other than Pa and M, like bedaquiline, helped by its high and long exposure in blood, may drive the efficacy of this powerful drug combination sufficiently to reduce effects of exposure differences for Pa and M, between the daily and weekend holiday schedules. Current PK analysis focuses on the fifth day of treatment (i.e. the end of the week), showing only minor differences in exposure on that day. It’s clear that exposure levels tend to decrease on the 5/7 treatment schedule after the weekend holiday period. However, this observation would require a more thorough investigation, likely through Population Pharmacokinetic (POPPK) modeling in the future, to provide more detailed insights.

Unlike CFU, which quantifies the bacteria that can grow to form a colony on a plate, RS ratio is a measure of ongoing rRNA synthesis, providing information on pathogen health and injury (i.e., RS ratio is high in actively replicating bacteria, and is greatly diminished in drug-injured or quiescent bacteria [23]. It is interesting to note that the two-day dosing holiday appears to significantly impact only the less effective 2RHZE/2RH drug combination, including drug with short half-life and where PK measures did not suggest varied exposures with or without a weekend dosing holiday. It is possible that more effective combination drug regimens like BPaMZ, which rapidly and profoundly reduce both lung CFU and RS ratio, are more forgiving owing to profound injury they cause to pathogen (i.e., PAE). Further studies to evaluate this proposal are warranted.

The data from the present study supports the importance of daily dosing of 2RHZE/2RH to optimize treatment efficacy. The forgiveness of BPaMZ with respect to treatment of TB patients is not yet known. However, these data demonstrate the potential utility of the BALB/c RMM, with CFU and RS ratio pharmacodynamic measures as a tool to evaluate regimen forgiveness for missed doses of new TB drug regimen combinations. Like all models, the BALB/c RMM has limitations, including pathology that does not include all lesion types seen in TB patients, such as those exhibiting caseous necrosis and cavitation. For this reason, the extent to which findings from the BALB/c RMM translate to the clinic is hard to predict, and it is likely that evaluation of forgiveness of regimens will require the development of additional tools, to generate data that may be used in conjunction with that from this mouse TB model. Further studies in the present model will better inform its contribution to an integrated approach to forgiveness evaluation.

Notwithstanding these limitations, the more limited impact of weekend dosing holidays on the derived T90 of BPaMZ suggests that forgiveness/impact of missed doses may be less relevant to clinical outcomes for drug combinations that include long half-life drugs (eg B) or those that promote profound pathogen injury (i.e., long PAE) than for RHZE/RH.

## MATERIAL AND METHODS

### Animals and ethics

All mouse experiments were carried out at the Evotec France SAS animal facility. This facility is accredited by the French Ministry of Agriculture and by the Association for Assessment and Accreditation of Laboratory Animal Care International (AAALAC). All studies were performed under the European Communities Council Directive (2010/063/EU) for the care and use of laboratory animals and approved by local Ethical Committee CEPAL: CE 029 and authorized by the French Ministry of Education, Advanced Studies and Research. Six weeks old female BALB/cJRj from Janvier Laboratories were used in these studies. Mice were group housed in bioconfined cages (Isocage, Tecniplast®) under a 12h light: 12h dark with free access to filtered water and a standard rodent diet (AO4C, Safe, France). An ambient temperature of 22 ± 2°C, a relative humidity of 55 ± 10 % and a negative pressure of −20Pa were maintained throughout the study. All mice were allowed to acclimatize to their new environment for at least 5 days.

### Drug Formulations and Dosing Strategies

Drugs were acquired and formulations prepared for dosing as follows: Bedaquiline (B, LTK Laboratories) was formulated in 20% 2-hydroxypropy-β-cyclodextrin for dosing at 25mg/kg; Pretomanid (Pa, Chemshuttle) was formulated in 10% Hydroxy-propyl-beta-cyclodextrin and 2% soy lecithin for dosing at 100mg/kg; Moxifloxacin (M, LTK Laboratories) and Pyrazinamide (Z, Sigma) were co-formulated with in water for dosing at 100mg/kg and 150 mg/kg respectively. Rifampicin (R, Sigma) was prepared in water for dosing at 10 mg/kg; Isoniazid (H, Sigma), Pyrazinamide (Z, Sigma) and Ethambutol (E, Sigma) were co-formulated in water for dosing at 10, 150 and 100mg/kg respectively. For the BPaMZ groups, B and Pa were individually administered then MZ co-formulated were dosed formulated. For RHZE/ RH groups, for the first 2 months, R was dosed individually, then spaced with HZE dosing which were co-formulated. For the following 2 months R and H were dosed individually. Mice were rested 2 h between doses.

### Relapsing Mouse Model

*Mycobacterium tuberculosis* H37Rv stock solution was prepared at exponential growth phase in 7H9 medium / 10% OADC (oleic acid-albumin dextrose-catalase) / 15% glycerol. At Day-14, mice were anaesthetized with 2.5% isoflurane in 97.5% oxygen and were intranasally infected with 50µL of *M*.*tb* H37Rv at an inoculum level of 4.5 Log10 CFU/mouse. Following the designated treatment period, mice were sacrificed 24h post last dosing. For relapse assessment, treatment was terminated at the end of each dosing period and lungs were collected 12 weeks after end of treatment (end of the relapse/cure phase). Lungs were collected and procedures conducted towards evaluation of endpoints as follows: For assessment of lung CFU and RS ratio, lung collection was performed as follows: The right upper lung lobes (1/3^rd^ of lung) were dissected and snap-frozen and kept at −80°C. The remaining left lung, lower right and accessory lobes (2/3^rd^ of lung) from this mouse were weighed and processed for bacteria enumeration. For assessment of relapse, collection of whole lungs from mice was performed after 12 weeks off treatment; whole lungs were harvested, weighed and processed for bacteria enumeration. In all cases, for bacterial enumeration, lung samples were homogenized and plated undiluted, or serial diluted on 7H11-OADC + 0.4% activated charcoal plates and incubated at 37°C for CFU quantification.

### PK Analysis

After 8 weeks of treatment, for each combination, (6 mice per group) blood was collected from the tail vein at 0.5, 2.5, 5.5, 7.5, 9 and 24 hours post-first compound dosing. At the terminal time point, blood was collected by cardiac puncture for plasma preparation and lungs were collected at 2.5h (3 mice) and 24h (3 mice) post first compound dosing. Plasma was prepared and stored at −80°C until analysis. Lungs were collected, weighed and stored at −80°C until processing for lung concentration analysis. After blood and lung processing, each compound, including bedaquiline M2 metabolite (BDQ-M2), was quantified by liquid chromatography tandem mass spectrometry (LC-MS/MS) methods.

Briefly, blood plasma or lung homogenates extracts were diluted with extracted control matrix as appropriate for the calibration range used. The supernatant (75 µL) was diluted 1:1 with 75 µL of water (Acetonitrile for Pyrazinamide, Isoniazid, Ethambutol and Rifampicin) prior to analysis. Bespoke chromatographic conditions were set up for: 1) BDQ, BDQ-M2, Pretomanid and Moxifloxacin, with: Phase A: Water with 0.1% formic Acid and Phase B: Acetonitrile with 0.1% formic Acid and 2) Pyrazinamide, Isoniazid, Rifampicin and Ethambutol with Phase A: Water with 0.5% formic Acid and Phase B: Acetonitrile with 0.5% formic Acid. Liquide Chromatographic gradients (phase A versus phase B) were applied for these 2 groups of compounds. The following mass spectrometry conditions were used for analysis of the compounds:

- Ionization mode : ESI positive mode
- Capillary Voltage : 3kV
- Desolvation Temperature : 600°C

The LC-MS parameters (transition) are reported in the (supplementary data). The limit of quantification (LOQ) of each compound is given in supplementary data.

### RS ratio® measurement

Lung samples that were collected and snap-frozen in Precellys tubes underwent 2-step homogenization and bead-beating, quantitative RNA extraction, fluorometric quantification, quality assessment via electrophoresis, reverse transcription, qPCR and digital PCR according to the standardized workflow. This workflow is accompanied with analysis of multiple process and procedural control specimens described by Walter et al [23]. The rRNA synthesis ratio (RS ratio®) was calculated as follow: unstable rRNA/stable rRNA =EST1/16S rRNA*104,000 [23].

### Logistic Emax Model

A Logistic Emax model was applied on data obtained from the current study plus historical ones, pooling together data of the same treatment collected from different studies (i.e. within the historical and current datasets). The historical dataset used consisted of data utilized in the development of the logistic regression model developed by Berg et al [22] in addition to historical data generated at Evotec.

The model used in this paper has the same structure as that described in Clary et al (to be submitted to AAC) [25], with the exception of the random effect on γ (i.e. η2) which is not considered in this work since its addition to the model did not provide significant improvement to model performance. For each regimen, γ and T50 parameters were estimated and subsequently used to calculate the related time to 90% cure (T90), according to the following formula:

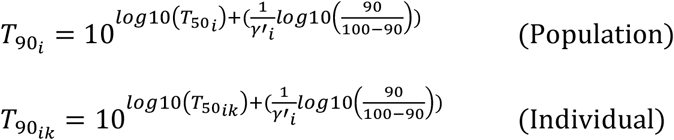

Regimens (in the historical dataset) that share the same drugs but were administered at different dose levels or with differing dose schedules were parametrized with different T50 parameters, while assuming the same γ value (i.e. steepness of the relapse-time curve). In this way it is possible to limit the number of parameters to be estimated and thus overcome identifiability issues.

Once all these values were computed for the combinations evaluated in this RMM study, a specific ranking of the two combinations for the two dosing schedules was drawn up in order to compare their performance.

The model was developed using NONMEM® 7.5.1 and data handling was performed through SAS® 9.4.

### Statistical analysis

CFU/lung was Log10 transformed before analysis. Zero CFU counts were replaced with 1 prior to transformation to allow calculation. The effect of dosing schedule 5/7 and 7/7 on number of CFU in lung and on RS ratio® results were evaluated for BPaMZ and 2RHZE/2RH considering the period of treatment using a two-way ANOVA followed by LSD Fisher test. The effect of the dosing schedule on derived T90 was compared using a Z-test. P-values <0.05 have been considered statistically significant.

## Acknowledgements

*“This work was supported by the Gates Foundation [BMGF01 INV-008993]. The conclusions and opinions expressed in this work are those of the author(s) alone and shall not be attributed to the Foundation. Under the grant conditions of the Foundation, a Creative Commons Attribution*

*4*.*0 License has already been assigned to the Author Accepted Manuscript version that might arise from this submission. Please note works submitted as a preprint have not undergone a peer review process*.*”*

*“We would like to thank the Evotec Translational Biology, BSL3 Infectious Diseases and BioServices and Welfare teams for their valuable technical support through this work to support this project”*.

*“We would like to thank the Evotec Pharmacokinetics team for their valuable support and collaboration throughout this work. In particular, we are especially grateful to Simone Zannoni and Chiara Roversi for their insightful input and contributions to the analytical aspects of this work, as well as their valuable contributions to the review and refinement of the manuscript*.*”*

## REFERENCES

1. World Health Organization. Global Tuberculosis Report 2023 WHO, Global Tuberculosis Programme (who.int).

2. Alipanah N, Jarlsberg L, Miller C, Linh N N, Falzon D, Jaramillo E, Nahid P. 2018. Adherence interventions and outcomes of tuberculosis treatment: A systematic review and meta-analysis of trials and observational studies. PLoS Med 15(7): e1002595.

3. Cadosch D, Abel Zur Wiesh P, Kouyos R, Bonhoeffer S. 2016. The Role of Adherence and Retreatment in De Novo Emergence of MDR-TB. PLoS Comput Biol 12(3): e1004749

4. Imperial MZ, Nahid P, Phillips PPJ, Davies GR, Fielding K, Hanna D, Hermann D, Wallis RS, Johnson JL, Lienhardt C, Savic RM. 2018. A patient-level pooled analysis of treatment-shortening regimens for drug-susceptible pulmonary tuberculosis. Nat Med 24(11):1708–1715.

5. Chang KC, Leung CC, Yew WW, Chan SL, Tam CM. 2006. Dosing schedules of 6-month regimens and relapse for pulmonary tuberculosis. Am J Respir Crit Care Med 174(10): 1153–8.

6. Vernon AA, Iademarco MF. 2004. In the treatment of tuberculosis, you get what you pay for. Am J Respir Crit Care Med 170(10): 1040–2.

7. Vashishtha R, Mohan K, Singh B, Devarapu SK, Sreenivas V, Ranjan S, Gupta D, Sinha S, Sharma SK. 2013. Efficacy and safety of thrice weekly DOTS in tuberculosis patients with and without HIV co-infection: an observational study. BMC Infect Dis. 13: 468

8. Stagg HR, Thompson JA, Lipman MCI, Sloan DJ, Flook M, Fielding KL. 2023. Forgiveness Is the Attribute of the Strong: Nonadherence and Regimen Shortening in Drug-sensitive Tuberculosis. Am J Respir Crit Care Med 207 (2):193–205.

9. Fox WS, Strydom N, Imperial MZ, Jarlsberg L, Savic RM. 2023. Examining nonadherence in the treatment of tuberculosis: The patterns that lead to failure. Br J Clin Pharmacol 89:1965–1977.

10. World Health Organization, Target regimen profiles for tuberculosis treatment, 2023 update: https://www.who.int/publications/i/item/9789240081512

11. MacKenzie FM & Gould IM. 1993. The post-antibiotic effect. J Antimicrob Chemother 32: 519–537.

12. Pai MP, Cottrell ML, Kashuba ADM, Bertino JS, Jr. 2015. Pharmacokinetics and Pharmacodynamics of Anti-infective Agents. Mandell, Douglas, and Bennett’s Principles and Practice of Infectious Diseases 252-262.e2

13. Cevik M, Thompson LC, Upton C, Rolla VC, Malahleha M, Mmbaga B, Ngubane N, Abu Bakar Z, Rassool M, Variava E, Dawson R, Staples S, Lalloo U, Louw C, Conradie F, Eristavi M, Samoilova A, Skornyakov SN, Ntinginya NE, Haraka F, Praygod G, Mayanja-Kizza H, Caoili J, Balanag V, Dalcolmo MP, McHugh T, Hunt R, Solanki P, Bateson A, Crook AM, Fabiane S, Timm J, Sun E, Spigelman M, Sloan DJ, Gillespie SH; SimpliciTB Consortium. 2024. Bedaquiline-pretomanid-moxifloxacin-pyrazinamide for drug-sensitive and drug-resistant pulmonary tuberculosis treatment: a phase 2c, open-label, multicentre, partially randomised controlled trial. Lancet Infect Dis 24(9):1003–1014.

14. Xu J, Li SY, Almeida DV, Tasneen R, Barnes-Boyle K, Converse PJ, Upton AM, Mdluli K, Fotouhi N, Nuermberger EL. 2019. Contribution of Pretomanid to Novel Regimens Containing Bedaquiline with either Linezolid or Moxifloxacin and Pyrazinamide in Murine Models of Tuberculosis. Antimicrob Agents Chemother 63(5). Pii: e00021–19.

15. Li SY, Irwin SM, Converse PJ, Mdluli KE, Lenaerts AJ, Nuermberger EL. 2015. Evaluation of Moxifloxacin-Containing Regimens in Pathologically Distinct Murine Tuberculosis Models. Antimicrobial Agents and Chemotherapy 59(7): 4026–4030.

16. Tasneen R. 2011. Sterilizing Activity of Novel TMC207- and PA-824-Containing Regimens in a Murine Model of Tuberculosis. Antimicrobial Agents and Chemotherapy 55(12): 5485–5492.

17. Li SY, Tasneen R, Tyagi S, Soni H, Converse PJ, Mdluli K, Nuermberger EL. 2017. Bactericidal and Sterilizing Activity of a Novel Regimen with Bedaquiline, Pretomanid, Moxifloxacin, and Pyrazinamide in a Murine Model of Tuberculosis. Antimicrob Agents Chemother 61(9): e00913–17.

18. Zhang M, Li SY, Rosenthal IM, Almeida DV, Ahmad Z, Converse PJ, Peloquin CA, Nuermberger EL, Grosset JH. 2011. Treatment of tuberculosis with rifamycin-containing regimens in immune-deficient mice. Am J Respir Crit Care Med. 183(9):1254–61.

19. Lenaerts AJ, Chapman PL, Orme IM. 2004. Limitations to the Cornell model of latent tuberculosis infection for the study of relapse rates. Statistical Tuberculosis (Edinb) 84:361–364.

20. Mourik BC, Svensson RJ, de Knegt GJ, Bax HI, Verbon A, Simonsson USH, de Steenwinkel JEM. 2018. Improving treatment outcome assessment in a mouse tuberculosis model. Nature/Scientific report 8:5714.

21. Mudde SE, Ayoun Alsoud R, van der Meijden A, Upton AM, Lotlikar MU, Simonsson USH, Bax HI, de Steenwinkel JEM. 2022. Predictive Modeling to Study the Treatment-Shortening Potential of Novel Tuberculosis Drug Regimens, Toward Bundling of Preclinical Data. J Infect Dis 225(11):1876–1885.

22. Berg A, Clary J, Hanna D, Nuermberger E, Lenaerts A, Ammerman N, Ramey M, Hartley D, Hermann D. 2022. Model-Based Meta-Analysis of Relapsing Mouse Model Studies from the Critical Path to Tuberculosis Drug Regimens Initiative Database. Antimicro Agents Chemothera 66 (3): e01793–21

23. Walter ND, Born SEM, Robertson GT, Reichlen M, Dide-Agossou C, Ektnitphong VA, Rossmassler K, Ramey ME, Bauman AA, Ozols V, Bearrows SC, Schoolnik G, Dolganov G, Garcia B, Musisi E, Worodria W, Huang L, Davis JL, Nguyen NV, Nguyen HV, Nguyen ATV, Phan H, Wilusz C, Podell BK, Sanoussi ND, de Jong BC, Merle CS, Affolabi D, McIlleron H, Garcia-Cremades M, Maidji E, Eshun-Wilson F, Aguilar-Rodriguez B, Karthikeyan D, Mdluli K, Bansbach C, Lenaerts AJ, Savic RM, Nahid P, Vásquez JJ, Voskuil MI. 2021. Mycobacterium tuberculosis precursor rRNA indicates treatment-shortening activity of drugs and regimens. Nature Communications 12(1):2899.

24. Dide-Agossou C, Bauman AA, Ramey ME, Rossmassler K, Al Mubarak R, Pauly S, Voskuil MI, Garcia-Cremades M, Savic RM, Nahid P, Moore CM, Tasneen R, Nuermberger EL, Robertson GT, Walter ND. 2022. Combination of Mycobacterium tuberculosis RS Ratio and CFU Improves the Ability of Murine Efficacy Experiments to Distinguish between Drug Treatments. Antimicrob Agents Chemother 66(4): e0231021

25. Musisi E, Dide-Agossou C, Al Mubarak R, Rossmassler K, Ssesolo AW, Kaswabuli S, Byanyima P, Sanyu I, Zawedde J, Worodria W, Voskuil MI, Savic RM, Nahid P, Davis JL, Huang L, Moore CM, Walter ND. 2021. Reproducibility of the Ribosomal RNA Synthesis Ratio in Sputum and Association with Markers of Mycobacterium tuberculosis. Microbiol Spectr 9(2):e0048121

26. Clary J, Roberts JK, Hanna D, Tagliavini A, Sordello S, Upton A, Hermann D and Berg Al. to be submitted 2025. A Stochastic Simulation-Based Approach to Inform Relapsing Mouse Model (RMM) Study Design. To be submitted AAC.

27. Chen C, Ortega F, Alameda L, Ferrer S and Simonsson USH. 2016. Population pharmacokinetics, optimised design and sample size determination for rifampicin, isoniazid, ethambutol and pyrazinamide in the mouse. Eur. J. Pharm Sciences 93: 319–333.

28. Rouan MC, Lounis N, Gevers T, Dillen L, Gilissen R, Raoof A, Andries K. Pharmacokinetics and Pharmacodynamics of TMC207 and its N-desmethyl metabolite in a murine model of tuberculosis. Antimicrob Agents Chemother. 2012 Mar;56(3):1444–51.

29. Irwin SM, Prideaux B, Lyon ER, Zimmerman MD, Brooks EJ, Schrupp CA, Chen C, Reichlen MJ, Asay BC, Voskuil MI, Nuermberger EL, Andries K, Lyons MA, Dartois V, Lenaerts AJ. 2016. Bedaquiline and Pyrazinamide Treatment Responses Are Affected by Pulmonary Lesion Heterogeneity in Mycobacterium tuberculosis Infected C3HeB/FeJ Mice. ACS Infect Dis 2(4):251–267

30. Ahmad Z, Peloquin CA Singh R P, Drendorf S T, Ginsberg A, Grosset J H, and Nuremberg E. 2011. PA-824 exhibits Time-dependent activity in a murine model of Tuberculosis. Antimicrob Agents Chemother. 55(1): 239–245.

31. Siefert HM, Domdey-Bette A, Henninger H, Hucke F, Kohlsdorfer C. and Stass HH. 1999. Pharmacokinetics of the 8-methoxyquinolone, moxifloxacin: a comparison in human and other mammalian species. Journal of Antimicrobial Chemotherapy. 43, Suppl. B: 69–76.

32. Hendrix J, Mubarak RA, Bateman A, Massoudi LM, Rossmassler K, Kaya F, Zimmerman MD, Wynn EA, Voskuil MI, Robertson GT, Moore CM, Walter ND. 2025. Physiologic recovery of Mycobacterium tuberculosis from drug injury: A molecular study of post antibiotic effect in mice. BioRxiv preprint.

33. Wynn EA, Dide-Agossou C, Mubarak RA, Rossmassler K, Hendrix J, Voskuil MI, Obregón-Henao A, Lyons MA, Robertson GT, Moore CM, Walter ND. 2025. Deconvoluting drug interactions based on M. tuberculosis physiologic processes: Transcriptional disaggregation of the BPaL regimen in vivo. BioRxiv preprint.

34. Bustion AE, Ernest JP, Kaya F, Silva C, Sarathy J, Blanc L, Imperial M, Gengenbacher M, Xie M, Zimmerman M, Robertson GT, Weiner D, Via LE, Barry CE, Savic RM, Dartois V. 2025. The kinetics of bedaquiline diffusion in tuberculous cavities open a window for emergence of resistance. J Infect Dis. Jun Online ahead of print

