## supplemental Table 1 for "A modeling-based framework to evaluate forgiveness of TB drug combinations in a BALB/c relapsing mouse model"

Supplementary data :

| Compound | Transition | CV[1] | CE[2] |
|---|---|---|---|
| Moxifloxacin | 402.18 > 110.08 | 22 | 34 |
| Bedaquiline | 557.16 > 229.15 | 50 | 22 |
| Pretomanid | 360.06 > 175.09 | 16 | 28 |
| Pyrazinamide | 123.98 > 80.99 | 50 | 16 |
| Rifampicin | 823.44 > 95.02 | 16 | 50 |
| Isoniazid | 138.01 > 78.99 | 22 | 22 |
| Ethambutol | 205.18 > 116.03 | 16 | 16 |
| BDQ-M2 | 541.10 > 480.11 | 34 | 22 |

**Table S1: Liquid Chromatography –Mass Spectrometry (i.e. LC-MS) parameters**

[1] CV: Cone Voltage
[2] CE: Collision Energy

| Compound | LOQ | | |
|---|---|---|---|
| | Plasma (ng/mL) | Blood (ng/mL) | Lung (ng/g) |
| Bedaquiline | 6 | 6 | 250 |
| Pretomanid | 12 | 12 | 5 |
| Moxifloxicin | 6 | 6 | 5 |
| Pyrazinamide | 200 | 200 | 10 |
| BDQ-M2 | 3 | 60 | 38500 |
| Rifampicin | 30 | 40 | 250 |
| Isoniazid | 40 | 300 | 250 |
| Pyrazinamide | 400 | 2000 | 200 |
| Ethambutol | 10 | 100 | 25 |

**Table S2: LIMIT OF QUANTIFICATION (LOQ)**

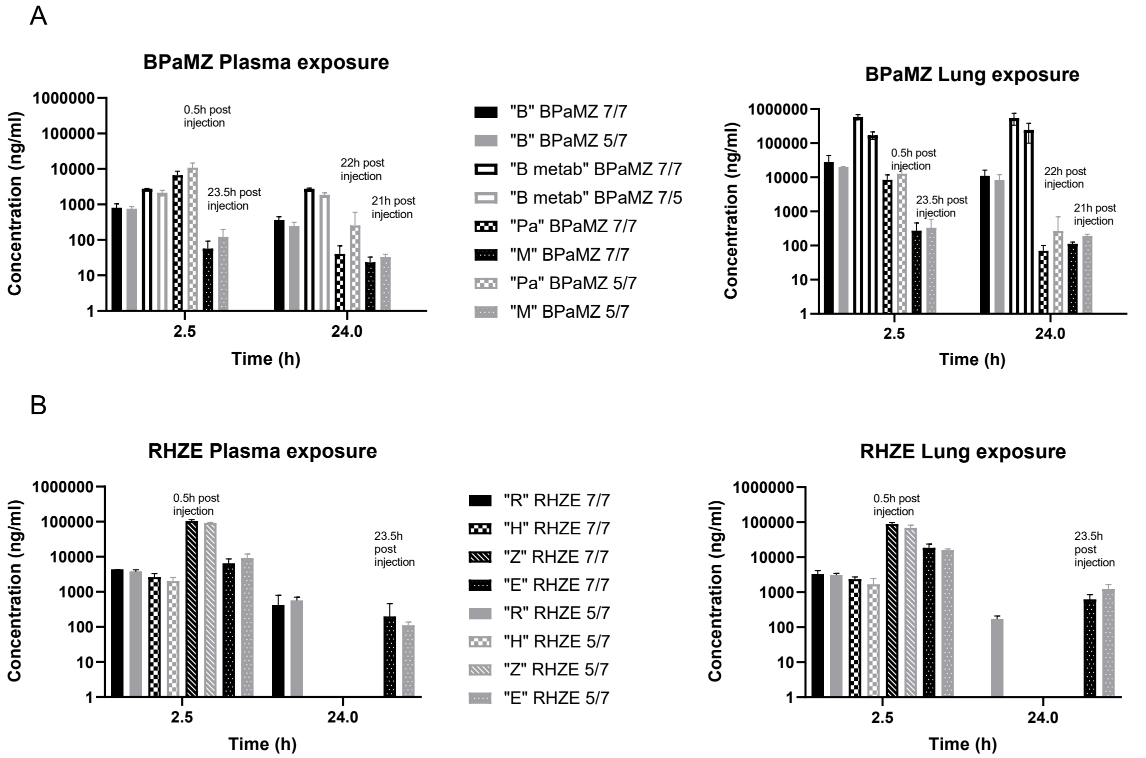


**Figure S3 Plasma and Lung exposure after 8 weeks of treatment the fifth day of the last week of treatment at 2.5 and 24h post dosing**

After 8 weeks of treatment, for each combination, at terminal time point, blood was collected by cardiac puncture for plasma preparation and lungs were collected at 2.5h and 24h post first compound dosing. Individual drug exposure was determined in BPaMZ (A) and RHZE (B) after 5/7 or 7/7 dosing.
